# The linear and non-linear effects of CYP2C19 metaboliser status on DNA methylation: a methylome-wide association study

**DOI:** 10.1101/2023.09.07.555071

**Authors:** Chen Shen, Mark J Adams, Eleanor Davyson, Matthew Iveson, Andrew M McIntosh, Xueyi Shen

## Abstract

CYP2C19 metabolises many medications e.g., antidepressants, proton pump inhibitors and its enzymatic activity can be inferred from genetic variants within its encoding gene, linking to the efficacy of drug treatments and their side effects. It is however unclear if enzymatic activity is associated with local or widespread differences in DNA methylation. DNA methylation differences associated with CYP2C19 metabolising status may also have the potential to reveal interacting genes and pathways that underly CYP2C19 effects on drug response and health consequences. A methylome-wide association study was conducted in the Generation Scotland cohort (n=18,396) to investigate the linear and non-linear effects of CYP2C19 metaboliser status on genome-wide DNA methylation. Pathway enrichment analysis was conducted for the cytosine-guanine dinucleotide (CpG) probes significantly associated with CYP2C19 metaboliser status. We examined whether the associations between CYP2C19 metaboliser status and DNA methylation were independent of the⍰use of drugs that are inducers, inhibitors, or substrates of the CYP2C19 enzyme by interaction analysis. Forty-eight CpG probes were significantly associated with both linear and quadratic terms of CYP2C19 metaboliser status (P_Bonferroni_ < 0.05). These CpG are annotated to genes involving drug metabolism, inflammation, lipid level, and Type 2 diabetes. Pathway enrichment analysis showed enrichment in biological processes involving metabolic activities and the Cytochrome P450 pathway. However, DNA methylation signals associated with CYP2C19 metaboliser status⍰did not vary by CYP2C19-related medication use. This research suggests that genetically-determined CYP2C19 metaboliser status is associated with both local and distal DNA methylation. These associations are independent of whether individuals were receiving drugs that are related to this enzyme.

## Introduction

The activity of many drug-metabolising enzymes influences drug response and side effects. Therefore, drug metabolising status is likely to have an important role in precision medicine by guiding the selection and dosage of medications for patients. The cytochromes P450 (CYP) superfamily comprises multiple genes that code for enzymes involving the metabolism of various exogenous and endogenous compounds such as prescribed drugs, environmental pollutants, cholesterol, steroids, and Vitamin D ^1^. The CYP family 2 subfamily C, polypeptide 19 (CYP2C19) enzyme, encoded by *CYP2C19* gene on chromosome 10 (10q24) ^2^, plays a key role in the metabolism of a wide range of clinically relevant drugs such as antidepressants, proton pump inhibitors (PPI), and antiplatelet medicine ^3-5^.

Genetic variants within the *CYP2C19* gene (e.g., *CYP2C19*x2, *CYP2C19*x3) are associated with reduced or loss of CYP2C19-mediated drug metabolising activity ^6^. Carriers of these variants have lower levels of active metabolites converted by these drugs, resulting in reduced response to treatment and/or increased side effects. For instance, evidence has indicated a relationship between *CYP2C19* genetic variants associated with poor metabolism and intolerance of citalopram, accompanied by discontinuation of treatment due to side effects ^7, 8^. Carriers of other *CYP2C19* variant alleles (e.g., *CYP2C19*x17, associated with ultrarapid metabolism) have increased drug metabolism capacity, which may result in a lack of response to standard antidepressant and PPI treatments due to an unusually rapid clearance of drugs ^8, 9^. This may indicate a potential non-linear relationship between CYP2C19 metaboliser status and health implications with drug use. It has been found that CYP2C19 metaboliser status has a non-linear relationship with antidepressant switching ^10^. *CYP2C19* polymorphism is also related to the metabolism of wider environmental exposures such as toxicants (e.g., heavy metals, organic pollutants) and nutrients (e.g., omega-3 fatty acids) ^11^. It has also been shown that individuals with high CYP2C19 enzymatic capacity (i.e., rapid/ultrarapid metaboliser) had more depressive symptoms regardless of antidepressant treatment ^12, 13^, but the biological mechanism is still unknown.

Epigenetic change can act as an environmental archive of exogenous exposures (including medication use) by reflecting duration, intensity, and individual susceptibility to adverse environmental risk factors ^14^. DNA methylation at cytosine-phosphate-guanine (CpG) sites is one of the most commonly investigated epigenetic processes. DNA methylation has established connections with environmental factors and provides important insights into the functional genomic mechanisms underlying common diseases ^15^. Previous methylome-wide association studies (MWAS) have found DNA methylation at specific CpG sites in relation to major depressive disorder and antidepressant use ^16-18^. However, to our knowledge, no studies have investigated the association between metaboliser status determined by *CYP2C19* variants and DNA methylation. It is unclear if enzymatic activity is associated with local or widespread differences in DNA methylation, which may have the potential to reveal interacting genes and pathways that underly CYP2C19 effects on drug response and health consequences. Whilst linear trends can inform DNA methylation patterns in relation to CYP2C19 metaboliser status, it is possible that associations are driven by genetic determinants of DNA methylation (methylation quantitative trait loci (mQTL)). Investigation of the non-linear relationship can help understand how CYP2C19 metaboliser status affects DNA methylation, where current evidence is limited.

The present study investigated the methylome-wide association for CYP2C19-determined metaboliser status from a large cohort, Generation Scotland (GS). We investigated both linear and non-linear effects of CYP2C19 metaboliser status on DNA methylation. We also investigated whether the use of CYP2C19-related drugs interacted with CYP2C19 metaboliser status to affect DNA methylation.

## Methods

### Participants

GS is family-based, population study mainly consists of people with European ancestry from across Scotland. Details of this cohort have been reported elsewhere ^19^. Ethical approval for this study was obtained from the NHS Tayside Research Ethics Committee (05/s1401/89). Written consent was obtained from all participants.

### Genetic data

Genotyping was conducted on blood samples collected at baseline using IlluminaHumanOmniExpressExome-8v1.0 BeadChip (48.8%) or Illumina HumanOmniExpressExome-8 v1.2 BeadChip (51.2%) ^20^. Quality check is consistent with previous publications ^21^, which includes sex mismatch detection, population outlier removal, genotype quality control with a call rate < 98%, individual removal based on a missing rate > 2%, Hardy-Weinberg equilibrium P value < 1×10^-6^, INFO score > 0.1, and MAF⍰≤ ⍰1%. Imputation in GS was conducted using the Sanger Imputation Server with the HRC v1.1 as the reference sample ^22^.

### CYP2C19 metaboliser status

CYP2C19 metaboliser status was derived using the PGxPOP pipeline to assign likely metaboliser status based on diplotypes ^23^. Five levels were included: poor metaboliser, intermediate metaboliser, normal metaboliser, rapid metaboliser, and ultrarapid metaboliser.

The *CYP2C19* gene is highly polymorphic with 38 variant alleles that encode CYP2C19 enzyme with no function (e.g., *CYP2C19*\*2), decreased function (e.g., *CYP2C19*\*9), normal function (e.g., *CYP2C19*\*1), or increased function (e.g., *CYP2C19*\*17). We ran PGxPOP on phased, imputed genotypes for chromosome 10 with hg19 coordinates and extracted phenotypes for the *CYP2C19* gene. Variants used to assign haplotypes were rs4244285 (*CYP2C19*\*2), rs4986893 (*CYP2C19*\*3), and rs12248560 (*CYP2C19*\*17), with CYP2C19*1 as the reference. According to the diplotype to phenotype definitions, participants with two no-function alleles were categorised as “poor metabolisers” (e.g., *CYP2C19*\*2/*2). Participants with two decreased function alleles or one normal/decreased function allele and one no-function allele were categorised as “intermediate metabolisers” (e.g., *CYP2C19*\*1/*2). Participants with two normal function alleles or one decreased function and one increased function alleles were categorised as “normal metabolisers” (e.g., *CYP2C19*\*1/*1. Participants with one increased function allele were categorised as “rapid metabolisers” (e.g., *CYP2C19*\*1/*17). Participants with two increased function alleles were categorised as “ultrarapid metabolisers” (e.g., *CYP2C19*\*17/*17).

### DNA methylation data

DNA methylation data was extracted from blood stored at the baseline ^24^. The protocol of preprocessing and quality check has been described in detail elsewhere ^24^. In short, CpG sites were removed if they had an outlying beta value (>+/-3 SD, P < 0.001), detection P value <0.005, and low or outlying bead counts (< 3 or >5% of the entire sample). Participants with sex mismatch with self-reported data and an outlier detection P < 0.01 for more than 5% of all CpG sites were removed from the analysis. Cross-hybridising and polymorphic CpG sites mapping to common SNPs (minor allele frequency (MAF) > 0.05) were removed from the analysis^25^. M-values were used for further analyses ^19^. To account for relatedness, M-values were residualised against the genomic relationship matrix created using GCTA ^26^. The residualised M-values were then carried into MWAS ^27^.

### MWAS statistical model

MWAS was conducted using the Omic-data-based complex trait analysis (OSCA) software (version 2.0) ^28^. Results from the linear regression model were presented as the main findings. The mixed-linear-model-based method (MOA) was used as a secondary method, accounting for collinearity between CpG sites. First, we set residualised M-values as the independent variable and the linear term of CYP2C19 metaboliser status as the outcome variable (continuous variable). Covariates are age, sex, first ten genetic principal components (PCs), smoking status (current/past/non-smoker), pack-years, first 20 methylation PCs, DNA methylation batch, DNA methylation-estimated cell proportions (CD8+T, CD4+T, natural killer cells, B cells and granulocytes). Bonferroni correction was applied across the entire methylome, with a P value < 6.64 x10^-8^ (0.05/752,741) set as the significance threshold. Subsequently, we conducted another MWAS to investigate the non-linear relationship between CYP2C19 metaboliser status and DNA methylation. We created the quadratic term of CYP2C19 metaboliser status as the outcome variable and adjusted for the same covariates as above plus the linear term of CYP2C19 metaboliser status. Bonferroni correction was also applied across the entire methylome.

### Pathway enrichment analysis

We used the ‘gometh’ function from the ‘missmethyl’ R package for Gene Ontology (GO) and Kyoto Encyclopedia of Genes and Genomes (KEGG) ^29^. Gometh analysis accounts for the number of CpG sites per gene to balance the statistical bias introduced by differing number of probes per gene present on the array, and CpGs that are annotated to multiple genes ^30^. Significant CpG sites found in the above two MWAS were used as the target list and the entire EPIC array as the background list. GO terms and KEGG pathway enrichment analyses were conducted separately. Default settings were used for the analysis.

### Differentially methylated region (DMR) analysis

We conducted the DMR analysis using the ‘dmrff’ package in R ^31^. DNA methylation data from a randomly selected list of participants from GS (N=1,000) was used as the reference panel. Significant DMRs for linear and quadratic terms of CYP2C19 metaboliser status were identified based on the following criteria: (1) the distance between two nearby probes within a DMR was at most 500 base pairs; (2) false discovery rate (FDR)-adjusted P value <⍰0.05, and (3) the same direction of MWAS effect estimates for the individual probes within a DMR.

### Sensitivity analysis

#### Interaction effect of CYP2C19 metaboliser status and CYP2C19-related medication use on DNA methylation

We examined the interaction effect between CYP2C19 metaboliser status and CYP2C19-metabolised medications use on DNA methylation. This is to investigate whether the effect of CYP2C19 metaboliser status on DNA methylation is driven by medication use. A list of drugs and their relationship with CYP2C19 enzyme (inducer, inhibitor, and substrate) were extracted from Drugbank (Supplementary file 1, also see URL: https://go.drugbank.com/bio_entities/BE0003536). Drugs associated with the *CYP2C19* gene were selected from the pharmacogenomics database. This list was used to flag records in community-dispensed prescription data (Prescribing Information System; covering 2009 to 2021, regardless of formulation, dose or prescribing duration ^32^. We identified users of CYP2C19-metabolised drugs by including any participant with a flagged record less than 6 months prior to baseline assessment (n=1460), and the remaining participants were treated as controls (n=4866).

We extracted M-values of CpG sites that showed non-linear associations with CYP2C19 metaboliser status as the dependent variables given that such a pattern may indicate pharmacological implications. We checked the significance of the interaction term of CYP2C19 metaboliser status (categorised as three groups: normal, rapid/intermediate, and ultrarapid/poor) and CYP2C19-related medication use (categorised as four groups: not current user (n=4866), users of drugs that CYP2C19 act as inducer (n=67), inhibitor (n=230), and substrate(n=1163)) in linear regression models, adjusting for age, sex, genotyping array, first ten genetic PCs, smoking status, pack-years, first 20 methylation PCs, DNA methylation batch, and DNA methylation-estimated cell proportions as covariates. CYP2C19 metaboliser status and CYP2C19-related medication use were included in the model as nominal variables.

### mQTL sensitivity analysis

Significant CpG associations with the linear term of CYP2C19 metaboliser status may be explained solely by the *cis*-mQTL effects. To investigate whether there is any secondary signal, we conducted another MWAS, adding the M-value of the top CpG site annotated to *CYP2C19* as an additional covariate. We also examined the correlations between significant CpG sites within the linkage disequilibrium (LD) block (r^2^ > 0.8) of the top mQTL of the top CpG site. This is to account for the possibility that nearby methylation sites are picking up signals from the same LD block.

## Results

### MWAS

A total of 18,396 GS participants with quality-controlled genetic and DNA methylation data were included in the MWAS (41.2% male, mean age = 47.5 years, standard deviation (SD) of age = 14.9 years, Table 1). The linear model MWAS showed that 48 CpG probes were significantly associated with the linear term of CYP2C19 metaboliser status with the Bonferroni threshold (P < 6.64 x10^-8^). Table 2 shows the top 10 CpG sites that were associated with the linear term of CYP2C19 metaboliser status. The Manhattan plot and QQ plot are shown in Figure 1 and Figure S1, respectively. The most significant CpG sites are annotated to *CYP2C19, NOC3L, PDLIM1, TBC1D12*, and *PLCE1* genes on Chromosome 10 (Table S1). Specifically, faster metaboliser status was associated with two hypermethylated CpG sites (cg02808805 and cg00051662) and one hypomethylated CpG site (cg20031717), all annotated to the *CYP2C19* gene. The MOA MWAS showed similar results to the linear model (Table S2). All the 48 CpG sites detected in the linear model MWAS remained significant.

**Table 1.** Demographic characteristics for GS individuals with quality-controlled genetic and DNA methylation data (n=18,396)

|  |  |
| --- | --- |
| **Age in years, mean (SD)** | 47.5 (14.9) |
| **Sex, n (%)** |  |
| Female | 10,823 (58.8) |
| Male | 7573 (41.2) |
| **Wave, n (%)** |  |
| 1 | 5086 (27.7) |
| 2 | 4447 (24.2) |
| 3 | 8863 (48.2) |
| **Smoking status, n (%)** |  |
| Current smoker | 3138 (17.1) |
| Former smoker (quitted within last 12 months) | 445 (2.4) |
| Former smoker (quitted more than 12 months ago) | 4727 (25.7) |
| Never smoker | 9404 (51.1) |
| Missing | 682 (3.7) |
| **Pack years, mean (SD)** | 7.31 (14.11) |
| **CYP2C19 metaboliser status, n (%)** |  |
| Poor | 399 (2.2) |
| Intermediate | 4754 (25.8) |
| Normal | 7395 (40.2) |
| Rapid | 4984 (27.1) |
| Ultrarapid | 864 (4.7) |

**Table 2.** Top 10 CpG sites that were associated with CYP2C19 metaboliser status.

| CYP2C19 metaboliser status | CpG | CHR | BP | B | SE | P | Nearest gene |
| --- | --- | --- | --- | --- | --- | --- | --- |
| Linear term | cg20031717 | 10 | 96523248 | -1.626 | 0.039 | $<2.22 \times 10^{-308}$ | *CYP2C19* |
| | cg02808805 | 10 | 96521820 | 1.231 | 0.031 | $<2.22 \times 10^{-308}$ | *CYP2C19* |
| | cg07889765 | 10 | 96123159 | -0.802 | 0.021 | $<2.22 \times 10^{-308}$ | *NOC3L* |
| | cg10751070 | 10 | 96143568 | 1.397 | 0.023 | $<2.22 \times 10^{-308}$ | |
| | cg11776334 | 10 | 96046836 | -1.515 | 0.041 | $7.06 \times 10^{-303}$ | *PLCE1* |
| | cg08923894 | 10 | 96123172 | -0.661 | 0.018 | $1.63 \times 10^{-287}$ | *NOC3L* |
| | cg23961982 | 10 | 96109291 | -1.230 | 0.036 | $1.10 \times 10^{-262}$ | *NOC3L* |
| | cg16964198 | 10 | 96199371 | -0.961 | 0.028 | $2.83 \times 10^{-253}$ | *TBC1D12* |
| | cg00051662 | 10 | 96521086 | 0.827 | 0.030 | $1.50 \times 10^{-170}$ | *CYP2C19* |
| | cg15851404 | 10 | 96643549 | -0.834 | 0.040 | $2.67 \times 10^{-96}$ | |
| Quadratic term | cg10751070 | 10 | 96143568 | 1.478 | 0.028 | $<2.22 \times 10^{-308}$ | |
| | cg16964198 | 10 | 96199371 | 2.325 | 0.028 | $<2.22 \times 10^{-308}$ | *TBC1D12* |
| | cg20031717 | 10 | 96523248 | 1.775 | 0.044 | $<2.22 \times 10^{-308}$ | *CYP2C19* |
| | cg08280358 | 10 | 96189867 | 0.919 | 0.036 | $1.04 \times 10^{-143}$ | *TBC1D12* |
| | cg08923894 | 10 | 96123172 | -0.478 | 0.021 | $1.47 \times 10^{-114}$ | *NOC3L* |
| | cg07889765 | 10 | 96123159 | -0.551 | 0.024 | $2.53 \times 10^{-114}$ | *NOC3L* |
| | cg08925046 | 10 | 97008920 | -1.420 | 0.065 | $2.97 \times 10^{-106}$ | *PDLIM1* |
| | cg14219693 | 10 | 96928076 | -1.400 | 0.065 | $1.56 \times 10^{-102}$ | |
| | cg11776334 | 10 | 96046836 | -0.906 | 0.047 | $6.35 \times 10^{-82}$ | *PLCE1* |
| | cg24087710 | 10 | 96928657 | -1.033 | 0.054 | $6.54 \times 10^{-81}$ | |

**Figure 1.**
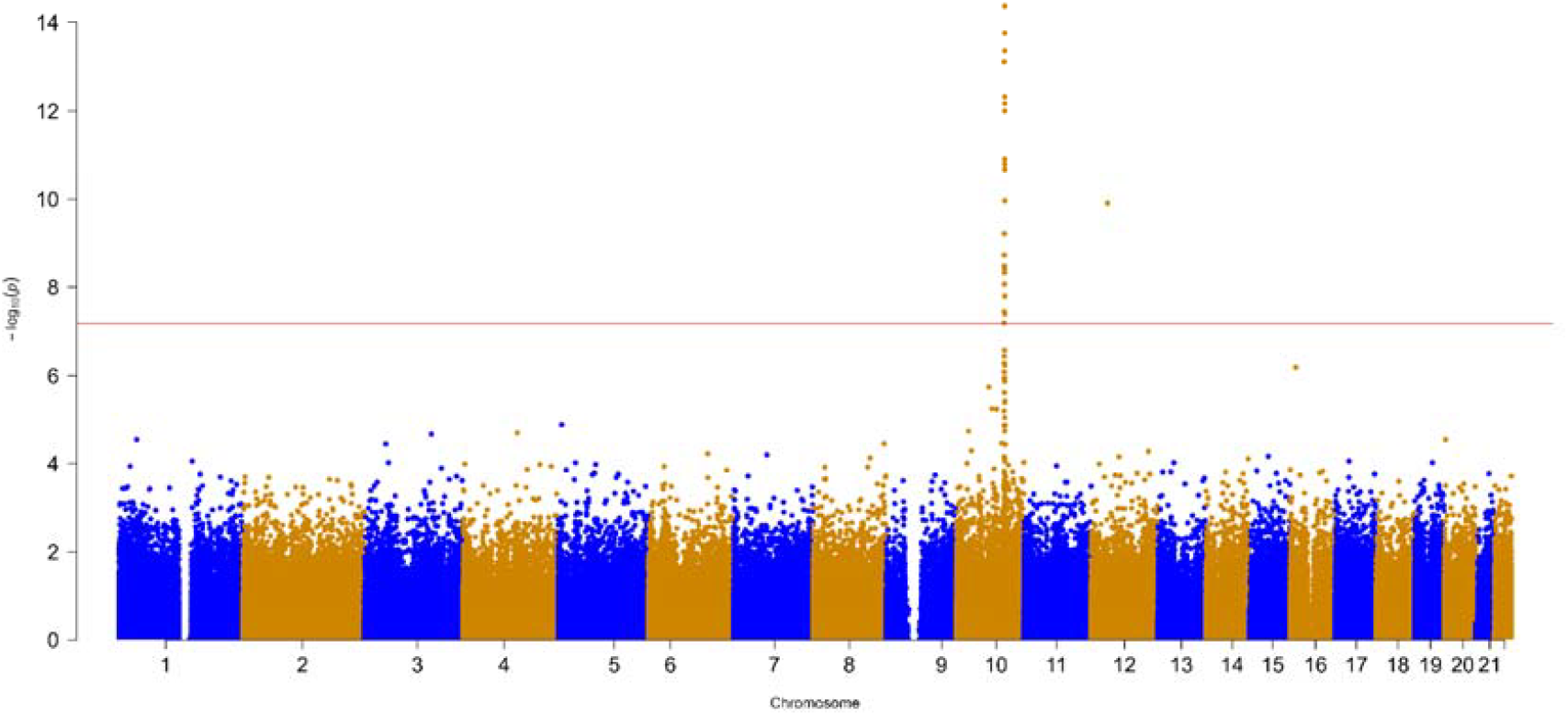
Manhattan plot for the MWAS of the linear term of CYP2C19 metaboliser status.

We found that 48 CpG sites were significantly associated with the quadratic term of CYP2C19 metaboliser status after Bonferroni correction (top 10 CpG sites shown in Table 2). The Manhattan plot and QQ plot are shown in Figure 2 and Figure S2, respectively. Whilst there were some overlaps of significant CpG sites with MWAS of the linear term of CYP2C19 metaboliser status, we found 19 CpG sites that showed non-linear associations with CYP2C19 metaboliser status (Figure 3, Table S3). DNA methylation of cg08280358, cg08883204, cg17725512, cg13512927, cg14302996, cg20426415, cg17014018, cg25674102, cg15160630, and cg04708601 showed a U-shaped relationship with CYP2C19 metaboliser status where normal metaboliser status had the lowest methylation level. DNA methylation of cg08925046, cg21800396, cg00087741, cg12575696, cg21636366, cg23210118, cg03013070, cg09847250, and cg14013452 showed an inverted U-shaped relationship with CYP2C19 metaboliser status where normal metaboliser status had the highest methylation level. These non-linear CpG sites are annotated to genes other than *CYP2C19* such as *TBC1D12, PDLIM1, ACSM6*, and *CYP2C18*.

**Figure 2.**
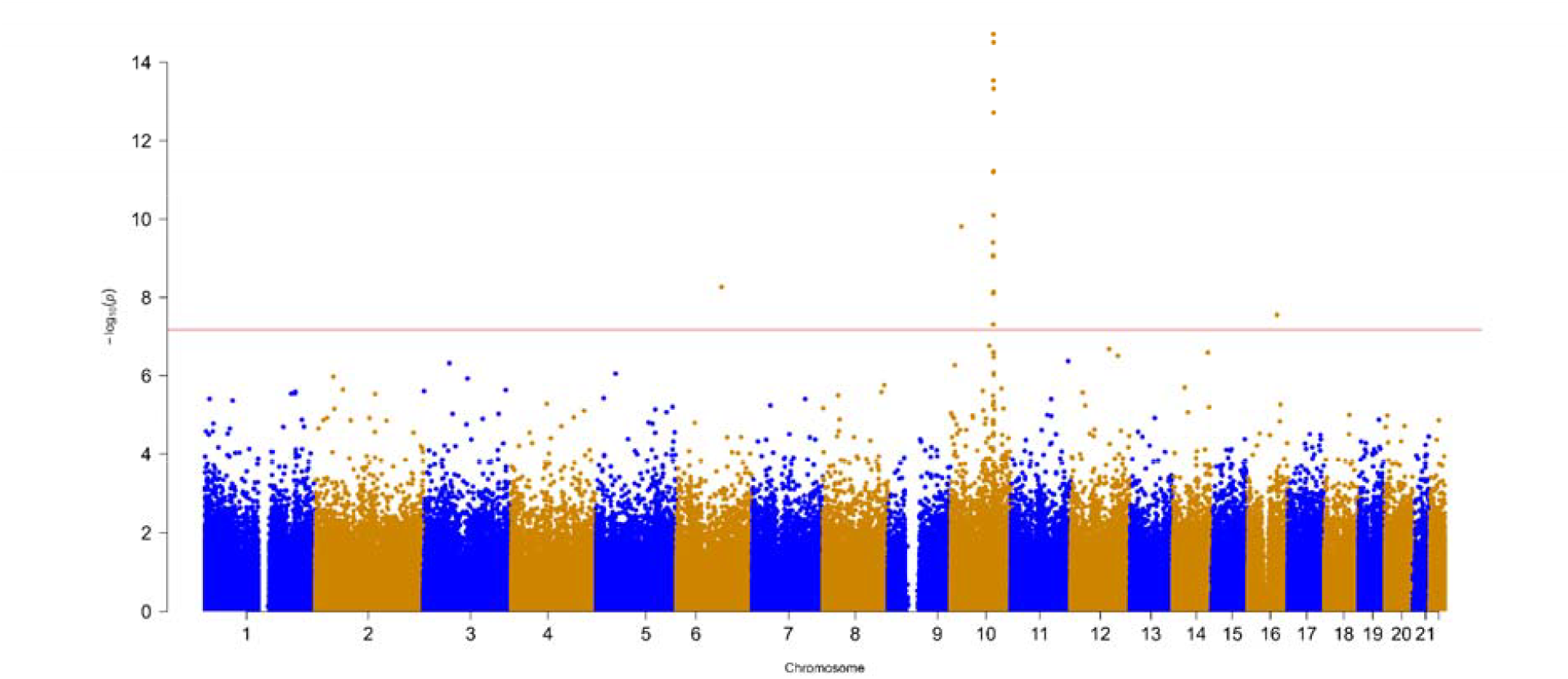
Manhattan plot for the MWAS of the quadratic term of CYP2C19 metaboliser status.

**Figure 3.**
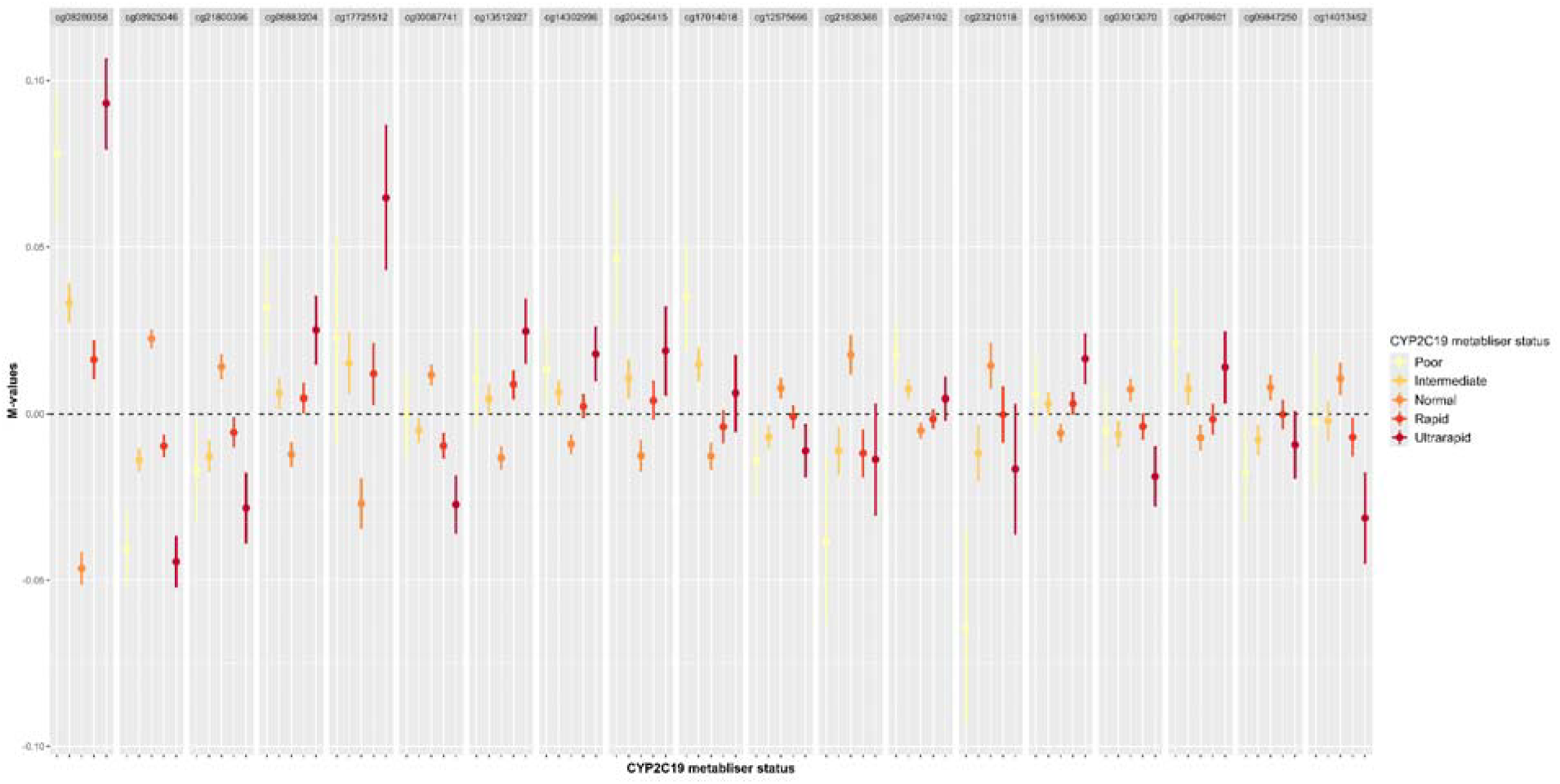
Non-linear effects of CYP2C19 metaboliser status on DNA methylation

### Pathway enrichment analysis

Analysis of the GO terms showed enrichment in biological processes involving metabolic activities and the P450 pathway. There were 162 and 206 enriched GO terms for MWAS of the linear term and quadratic term of CYP2C19 metaboliser status with a P value < 0.05, respectively. However, none was significant after the FDR correction. The top ten GO terms are listed in Table 3. Top pathways were driven by *CYP2C19, CYP2C18, PKP2*, and *PDLIM1* genes. The test of KEGG pathways showed that pathways such as chemical carcinogenesis and metabolic pathways, including drug metabolism, were enriched. There were 11 and 9 nominally significant KEGG pathways for MWAS of the linear term and quadratic term of CYP2C19 metaboliser status with a P value < 0.05, respectively. There was no significant pathway after FDR correction (Table 4). Nominally significant pathways were driven by *CYP2C19, CYP2C18, PKP2, PDLIM1, ACSM6, and PLCE1* genes.

**Table 3.**
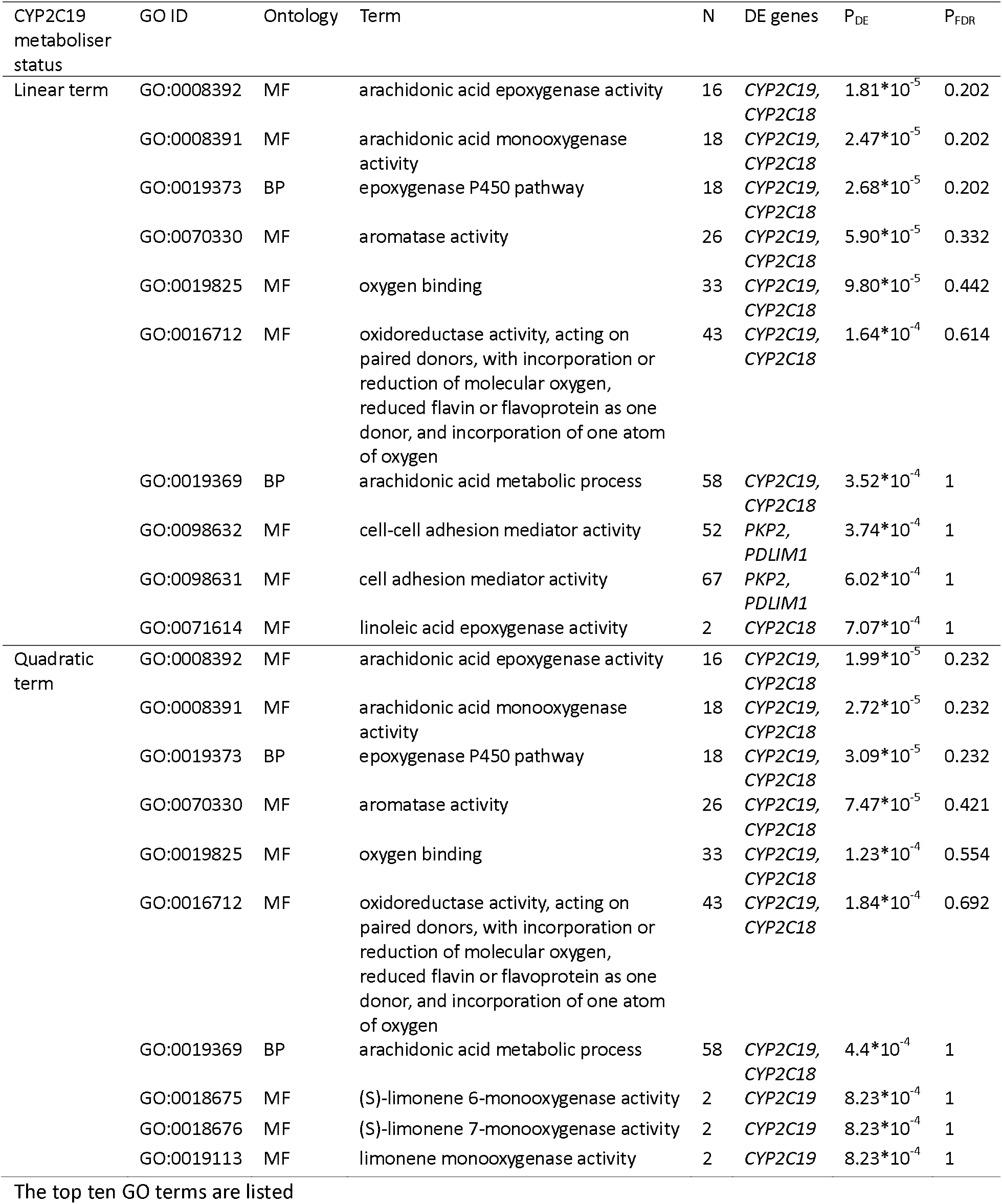
Results for gene ontology (GO) analysis for the MWAS on CYP2C19 metaboliser status at P threshold 6.64 x10^-8^. BP: Biological process; MF: molecular function; DE genes: genes that are differentially methylated.

**Table 4.** Kyoto Encyclopedia of Genes and Genomes (KEGG) pathway analysis for the MWAS on CYP2C19 metaboliser status at P threshold 6.64 x10^-8^. DE genes: genes that are differentially methylated.

| CYP2C19 metaboliser status | KEGG pathway ID | Description | N | DE genes | P<sub>DE</sub> | P<sub>FDR</sub> |
| --- | --- | --- | --- | --- | --- | --- |
| Linear term | hsa05204 | Chemical carcinogenesis - DNA adducts | 70 | *CYP2C19, CYP2C18* | 0.000 | 0.136 |
|  | hsa04726 | Serotonergic synapse | 113 | *CYP2C19, CYP2C18* | 0.002 | 0.279 |
|  | hsa01100 | Metabolic pathways | 1532 | *ACSM6, CYP2C19, CYP2C18, PLCE1* | 0.005 | 0.625 |
|  | hsa04820 | Cytoskeleton in muscle cells | 232 | *PKP2, PDLIM1* | 0.007 | 0.625 |
|  | hsa00650 | Butanoate metabolism | 27 | *ACSM6* | 0.013 | 0.809 |
|  | hsa00591 | Linoleic acid metabolism | 30 | *CYP2C19* | 0.013 | 0.809 |
|  | hsa00590 | Arachidonic acid metabolism | 61 | *CYP2C19* | 0.028 | 1 |
|  | hsa00830 | Retinol metabolism | 68 | *CYP2C18* | 0.029 | 1 |
|  | hsa00982 | Drug metabolism - cytochrome P450 | 71 | *CYP2C19* | 0.029 | 1 |
|  | hsa00562 | Inositol phosphate metabolism | 73 | *PLCE1* | 0.041 | 1 |
|  | hsa05412 | Arrhythmogenic right ventricular cardiomyopathy | 86 | *PKP2* | 0.048 | 1 |
| Quadratic term | hsa05204 | Chemical carcinogenesis - DNA adducts | 70 | *CYP2C19, CYP2C18* | 0.000 | 0.151 |
|  | hsa04726 | Serotonergic synapse | 113 | *CYP2C19, CYP2C18* | 0.003 | 0.501 |
|  | hsa01100 | Metabolic pathways | 1532 | *ACSM6, CYP2C19, CYP2C18, PLCE1* | 0.010 | 1 |
|  | hsa00650 | Butanoate metabolism | 27 | *ACSM6* | 0.015 | 1 |
|  | hsa00591 | Linoleic acid metabolism | 30 | *CYP2C19* | 0.016 | 1 |
|  | hsa00830 | Retinol metabolism | 68 | *CYP2C18* | 0.030 | 1 |
|  | hsa00982 | Drug metabolism - cytochrome P450 | 71 | *CYP2C19* | 0.030 | 1 |
|  | hsa00590 | Arachidonic acid metabolism | 61 | *CYP2C19* | 0.032 | 1 |
|  | hsa03320 | PPAR signaling pathway | 75 | *SORBS1* | 0.045 | 1 |

KEGG pathways with P<sub>DE</sub> < 0.05 are listed

### DMR analysis

A total of 5 DMRs annotated to *PDLIM1* (97051104 - 97051225; 97051255 - 97051319), *NOC3L* (96121776 - 96121853), *CYP2C18* (96442621 - 96443071), and an open-sea area (96928199 - 96928657) were identified as associated with the linear term of CYP2C19 metaboliser status at FDR-adjusted P value < 0.05 (Table 5). Faster metaboliser status was associated with two hypomethylated DMRs annotated to the *PDLIM1* gene, one hypermethylated DMR annotated to the *NOC3L* gene, and one hypomethylated DMR annotated to the *CYP2C18* gene. No significant DMR was identified in the *CYP2C19* gene. A total of 3 DMRs annotated to *NOC3L* (96123159 - 96123172), *PLCE1* (96048116 - 96048482), and *SORBS1* (97204891 - 97205147) were associated with the quadratic term of CYP2C19 metaboliser status at FDR-adjusted P value < 0.05.

**Table 5.** Significant differentially methylated regions identified from the MWAS analysis.

| CYP2C19<br>metaboliser<br>status | CHR | Start | End | Gene | Effect | SE | Adjusted P<br>values | CpGs |
| --- | --- | --- | --- | --- | --- | --- | --- | --- |
| Linear | 10 | 97051104 | 97051225 | *PDLIM1* | -1.234 | 0.141 | $1.39 \times 10^{-12}$ | cg11911874,<br>cg03178678 |
| | 10 | 97051255 | 97051319 | *PDLIM1* | -0.888 | 0.133 | $1.89 \times 10^{-05}$ | cg06542614,<br>cg05599883 |
| | 10 | 96121776 | 96121853 | *NOC3L* | 0.261 | 0.036 | $6.16 \times 10^{-07}$ | cg15209556,<br>cg17162216,<br>cg10164249 |
|  | 10 | 96442621 | 96443071 | *CYP2C18* | -0.660 | 0.116 | 0.01 | cg17014018,<br>cg04501839,<br>cg19852607 |
| | 10 | 96928199 | 96928657 | | 1.224 | 0.182 | $1.28 \times 10^{-05}$ | cg25102879,<br>cg24087710 |
| Quadratic | 10 | 96123159 | 96123172 | *NOC3L* | -1.357 | 0.061 | $1.50 \times 10^{-104}$ | cg07889765,<br>cg08923894 |
| | 10 | 96048116 | 96048482 | *PLCE1* | 2.592 | 0.369 | $1.62 \times 10^{-06}$ | cg15160630,<br>cg10680511 |
| | 10 | 97204891 | 97205147 | *SORBS1* | 1.726 | 0.204 | $2.08 \times 10^{-11}$ | cg03799335,<br>cg14302996 |

SE: standard error

### Sensitivity analysis

We investigated the interaction effect of CYP2C19 metaboliser status and CYP2C19-metabolised medication use on M-values of CpG sites that showed significant non-linear associations with CYP2C19 metaboliser status. No significant interaction effect was detected (P values for interaction term after FDR correction: all > 0.05, Table S4), indicating that the association between CYP2C19 metaboliser status and DNA methylation did not vary by CYP2C19-metabolised medication use.

When additionally adjusting for the M-value of the CpG that was most robustly associated with the linear term of CYP2C19 metaboliser status (cg20031717), the associations between other significant CpG probes (e.g., cg02808805, cg00051662) annotated to the *CYP2C19* gene and CYP2C19 metaboliser status were attenuated but still significant (Table S5), indicating that associations between CYP2C19 metaboliser status and DNA methylation were not explained solely from mQTL of the *CYP2C19* gene. The associations between significant CpG probes (e.g., cg07889765, cg11776334) annotated to other genes and CYP2C19 metaboliser status remained similar. We found 4 other CpG sites (cg00051662, cg02808805, cg14196507, cg15851404) within the LD block of the top mQTL (rs200889969) of cg20031717. Correlations between these CpG sites were low (coefficients ranged between −0.18 and 0.24, Figure S3).

## Discussion

We found that CYP2C19 metaboliser status had both linear and non-linear effects on DNA methylation in a large population health dataset. Significant CpGs were enriched in metabolic pathways involving P450 epoxygenase and drug metabolism, mostly driven by *CYP2C19* and *CYP2C18* genes. No interactive effect was observed between CYP2C19-related medication use and CYP2C19 metaboliser status on DNA methylation.

cg20031717 annotated to the *CYP2C19* gene was most robustly associated with the linear term of CYP2C19 metaboliser status. Other significant CpG sites annotated to the *CYP2C19* gene were cg02808805 and cg00051662, with hypermethylation within these two CpG sites found in faster metaboliser status. DNA methylation at both cg20031717 and cg00051662 has been reported to decrease with age but is not associated with any other traits ^33^. Hypermethylation within cg02808805 has been found associated with the prevalence of prostate cancer ^34^. This is corroborated by the KEGG analysis indicating enriched pathways involving chemical carcinogenesis.

Among the CpG sites showing non-linear relationships with CYP2C19 metaboliser status, hypermethylation of cg21800396 and cg00087741 and hypomethylation of cg08883204, cg14302996, cg25674102, and cg04708601 were observed in normal metaboliser status and are also associated with a higher level of C-reactive protein (CRP) and increased risk of Type 2 diabetes in other MWAS^34, 35^. Normal metaboliser status was also associated with hypermethylation of cg09847250 and hypomethylation of cg17014018 which are related to a lower level of CRP. However, neither of these two CpG sites is linked to Type 2 diabetes. These findings indicate that CYP2C19 metaboliser status may have implications for inflammation, and normal metaboliser status may link to increased risk of Type 2 diabetes.

Significant CpG sites and DMRs are mapped to multiple genes on Chromosome 10, such as *NOC3L, PDLIM1, PLCE1, CYP2C18, and SORBS1*. Of these, *SORBS1* is specifically relevant to non-linear CpG signal (cg14302996) and DMR. The protein encoded by this gene has functions in insulin signalling and stimulation, where dysfunction may be associated with insulin resistance and Type 2 diabetes ^36^. *NOC3L and PLCE1* are associated with low-density lipoprotein (LDL) cholesterol levels in the blood ^37^. Previous research highlights the important role CYP enzymes play in the metabolism of cholesterols ^38^. This explains that lipid profile, including LDL subfractions, predicts response to antidepressants, as found by another study ^39^. *PDLIM1* has been associated with metabolite levels of warfarin (an anticoagulant drug) and steroid hormone levels (biosynthesis promoted by CYP) ^40, 41^. *CYP2C18* is another CYP superfamily member associated with metabolite levels of warfarin and clopidogrel as well as serum metabolites ^40, 42, 43^.

Our sensitivity analysis did not detect the interaction effect between CYP2C19-related medication use and CYP2C19 metaboliser status on DNA methylation. This indicates that metabolites of these drugs and metabolic status determined by *CYP2C19* variants may affect DNA methylation independently. This is partly supported by previous MWAS in European ancestries on antidepressant use ^17, 18^. CpG signals associated with antidepressant use found in these two studies do not overlap with any significant CpG sites or DMRs in our study, indicating that CYP2C19 metaboliser status and antidepressant use may not share biological pathways. However, this has not formally been tested with other CYP2C19-metabolised drugs, e.g., warfarin, clopidogrel, to our knowledge.

Strengths of our study include that GS is the largest single sample MWAS to date, which allows adequate power to detect individual CpG sites and DMRs associated with CYP2C19 metaboliser status. Additionally, as CYP2C19-determined metaboliser status is genetically determined, reverse causality is not a concern. However, our study has some limitations. First, our analyses only included participants predominantly with European ancestry, so our findings may not be generalisable to individuals of other ancestries. Second, DNA methylation from the whole blood may not be representative of DNA methylation from the liver where the *CYP2C19* gene is mostly expressed and may obscure findings from individual cell types. There is a lack of evidence on the concordance of DNA methylation profile in relation to CYP enzymes between blood and liver tissues. Third, we categorised the current use of CYP2C19-metabolised medications as inducer, inhibitor, and substrate when examining its interaction with CYP2C19 metaboliser status affecting DNA methylation. Future study should gather more detailed information on medication use (e.g., dose, duration, side effect) and concentrations in blood to enable a better understanding of the biological effect of CYP2C19 metaboliser status together with CYP2C19-metabolised medication use on DNA methylation. Information on drug use is only available in a subset of participants who gave consent for data linkage to electronic health records, which might limit the power to detect a significant interaction effect. Finally, we did not replicate our analyses in another sample, which is required in future research to confirm our findings.

In conclusion, this MWAS in a large-scale cohort suggests that genetically-determined CYP2C19 metaboliser status is associated with both local and distal DNA methylation. CYP2C19 metaboliser status may have biological implications relevant to the response of certain drugs, inflammation, lipid level, and Type 2 diabetes. The use of CYP2C19-metabolised medications did not interact with CYP2C19 metaboliser status to affect DNA methylation, indicating that CYP2C19 metaboliser status may impact DNA methylation and metabolic pathways that do not depend on the use of medications metabolised by this enzyme. Future studies should confirm our findings in other populations and investigate how CYP2C19 metaboliser status interacts with detailed medication records on DNA methylation and health consequences.

## Data sharing

According to the terms of consent for GS participants, access to individual-level genetic, DNA methylation data and phenotypes need to be approved by the GS Access Committee (https://www.ed.ac.uk/generation-scotland/for-researchers/access). Application should be made to. Data dictionary for GS is available at https://datashare.ed.ac.uk/handle/10283/2988.

## Acknowledgements

This work is supported by the Wellcome Trust (220857/Z/20/Z, 104036/Z/14/Z and 216767/Z/19/Z) and the Medical Research Council (MR/W014386/1). For the purpose of open access, the author has applied a CC BY public copyright licence to any Author Accepted Manuscript version arising from this submission. In addition, DNA methylation profiling was supported by funding from NARSAD (27404) and the Royal College of Physicians of Edinburgh (SIM Fellowship). Genotyping of the GS samples was funded by the MRC and Wellcome Trust (104036/Z/14/Z). GS also receives support from the Chief Scientist Office of the Scottish Government Health Directorates (CZD/16/6) and the Scottish Funding Council (HR03006). This work has used the resources provided by the Edinburgh Compute and Data Facility (ECDF) (http://www.ecdf.ed.ac.uk/). CS was supported by the National Institute for Health Research (NIHR) Imperial Biomedical Research Centre (PSR917]. ED was supported by the UK Research and Innovation (EP/S02431X/1), UK Research and Innovation Centre for Doctoral Training in Biomedical AI at the University of Edinburgh, School of Informatics. The views expressed are those of the authors and not necessarily those of the sponsors.

We thank the participants and team members for their ongoing contribution to the recruitment, data management and technical and legal support for GS.

## Contributors

CS, AMM, and XS conceived the study concept. CS conducted MWAS data analysis and interpretation for GS. MJA, MI, and XS contributed to data preparation for GS. CS wrote the manuscript. MJA, ED, MI, AMM, and XS edited the manuscript. All authors read and approved the final manuscript.

