## Supplementary figures and tables for "The linear and non-linear effects of CYP2C19 metaboliser status on DNA methylation: a methylome-wide association study"

Figure S1 QQ plot for the MWAS of linear term of CYP2C19 metaboliser status.

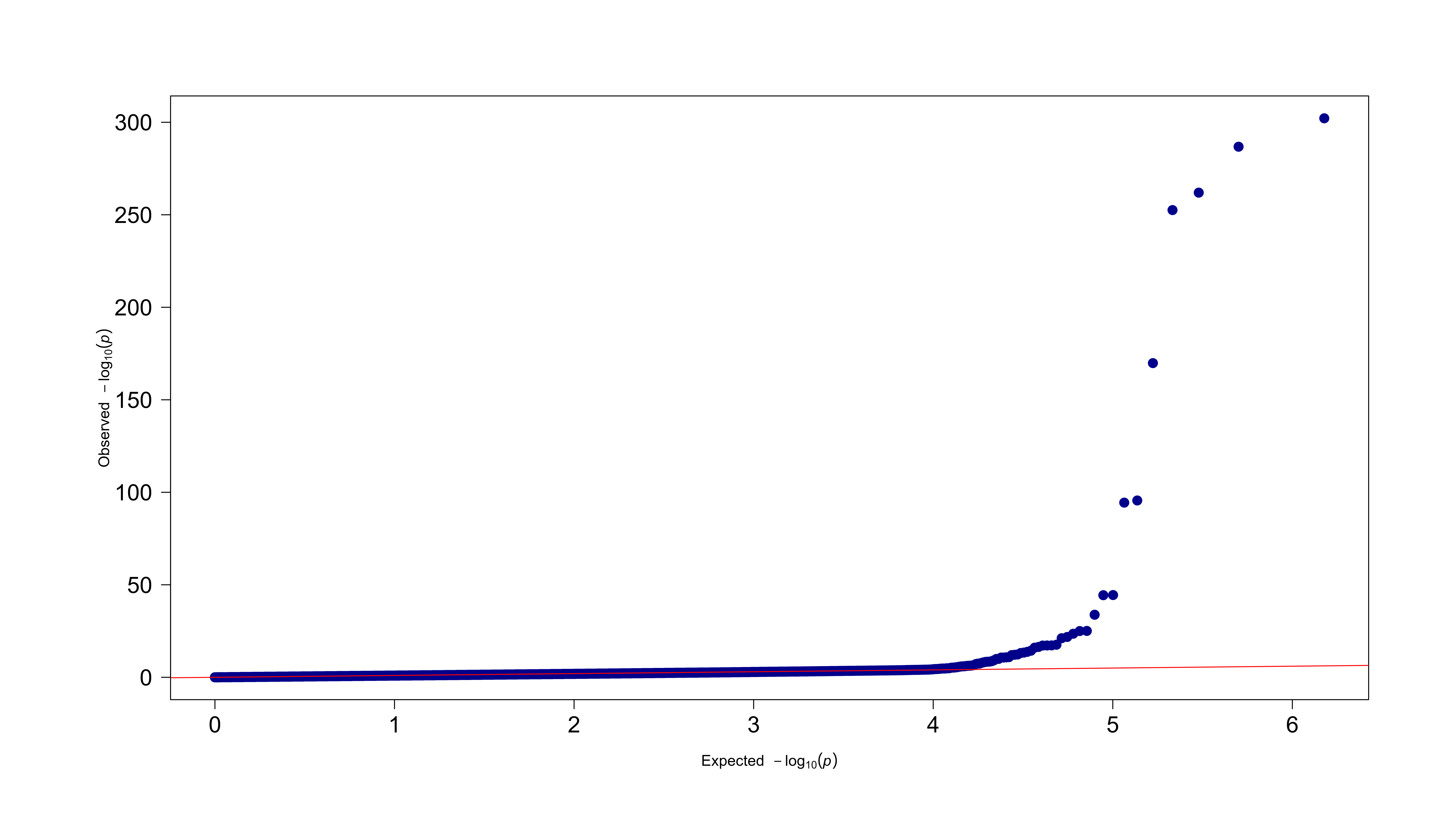

Figure S2 QQ plot for the MWAS of the quadratic term of CYP2C19 metaboliser status.

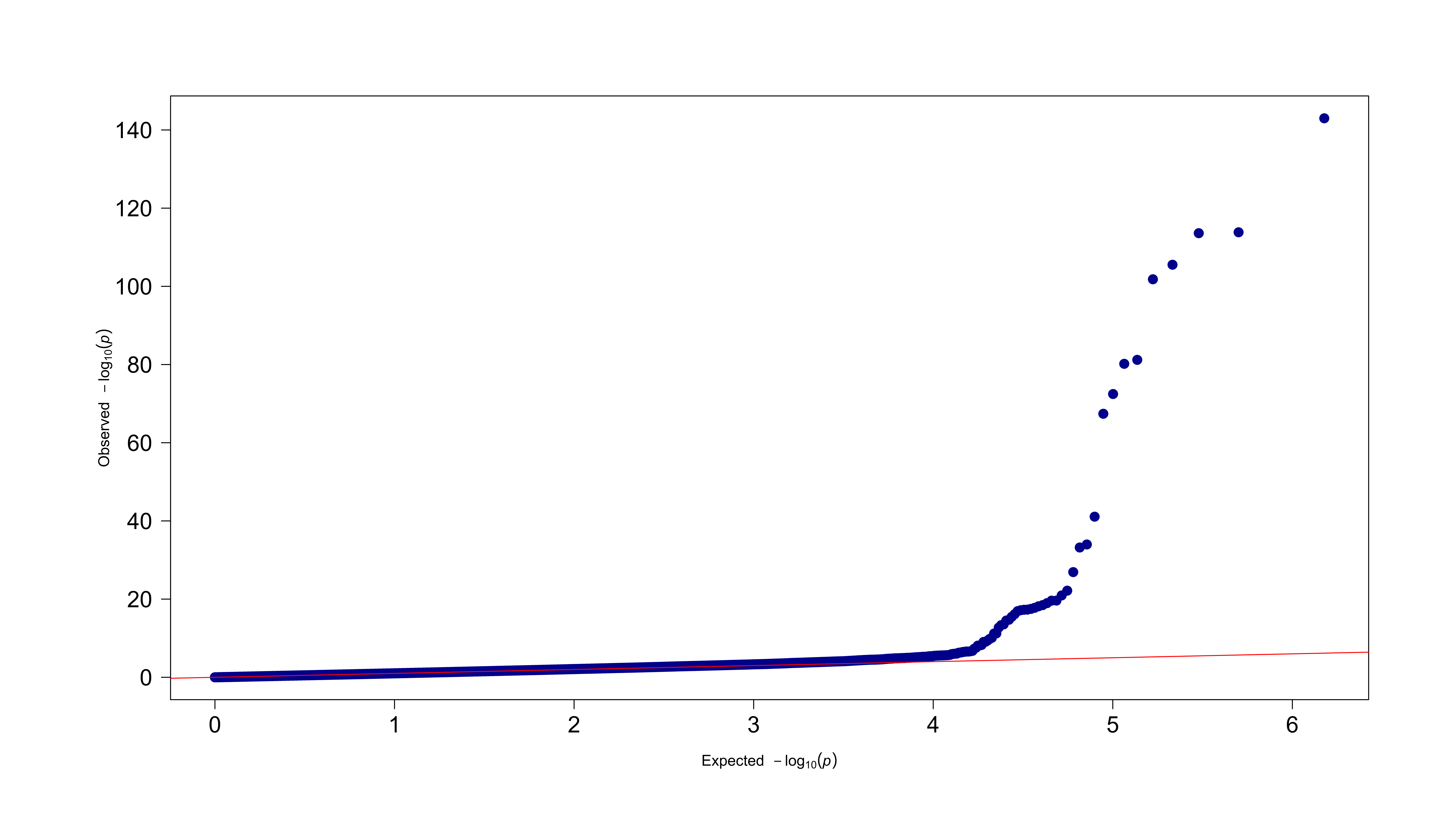

Figure S3 Correlation heatmap for CpG sites within the LD block of the nearest mQTL (rs12773342) to cg20031717

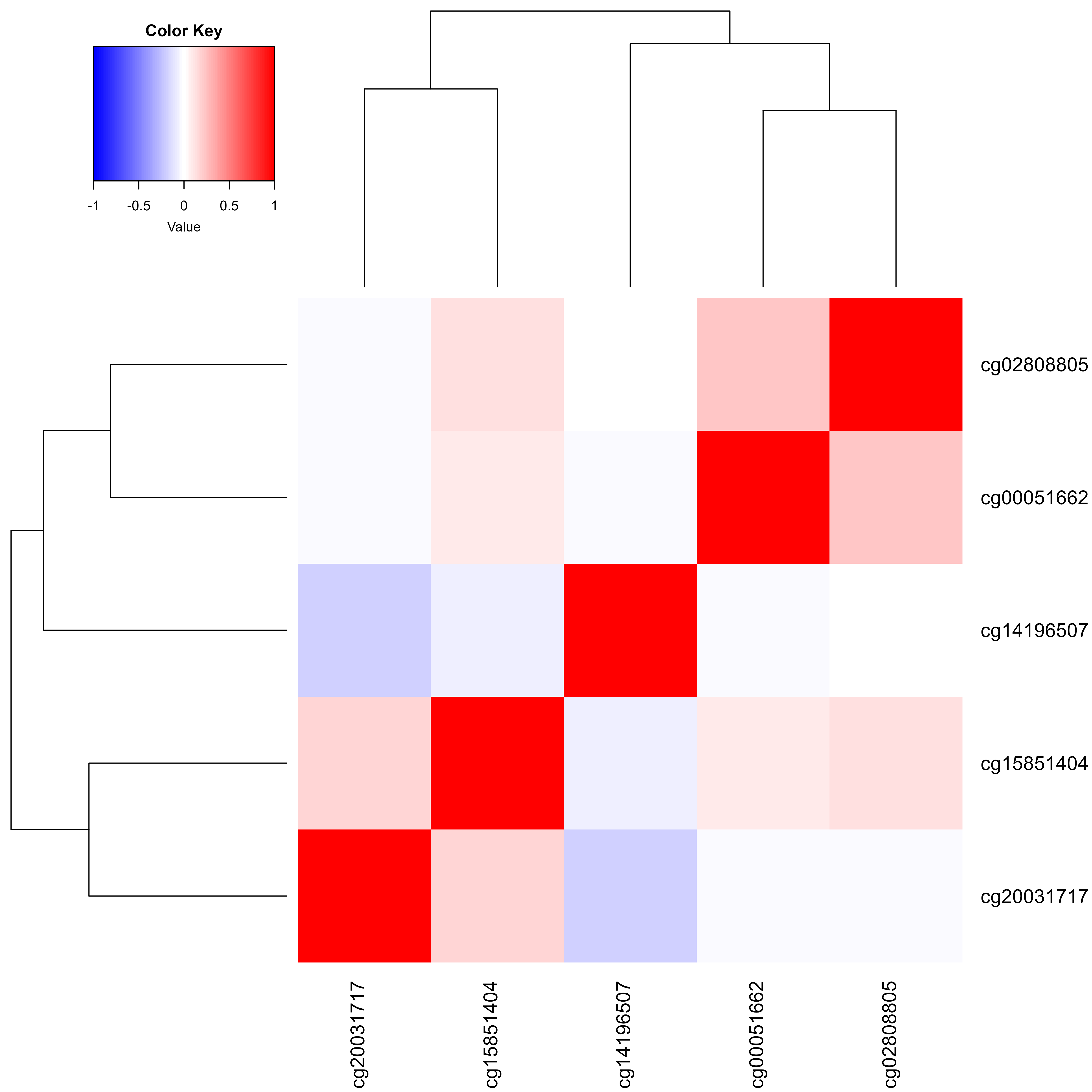

Table S1 Significant CpGs associated with the linear term of CYP2C19 metaboliser status (linear model MWAS analysis).

| CpG | CHR | BP | B | SE | P | Nearest gene |
| --- | --- | --- | --- | --- | --- | --- |
| cg20031717 | 10 | 96523248 | -1.626 | 0.039 | <2.22x10^-308^ | *CYP2C19* |
| cg02808805 | 10 | 96521820 | 1.231 | 0.031 | <2.22x10^-308^ | *CYP2C19* |
| cg07889765 | 10 | 96123159 | -0.802 | 0.021 | <2.22x10^-308^ | *NOC3L* |
| cg10751070 | 10 | 96143568 | 1.397 | 0.023 | <2.22x10^-308^ |  |
| cg11776334 | 10 | 96046836 | -1.515 | 0.041 | 7.06x10^-303^ | *PLCE1* |
| cg08923894 | 10 | 96123172 | -0.661 | 0.018 | 1.63x10^-287^ | *NOC3L* |
| cg23961982 | 10 | 96109291 | -1.230 | 0.036 | 1.10x10^-262^ | *NOC3L* |
| cg16964198 | 10 | 96199371 | -0.961 | 0.028 | 2.83x10^-253^ | *TBC1D12* |
| cg00051662 | 10 | 96521086 | 0.827 | 0.030 | 1.50x10^-170^ | *CYP2C19* |
| cg15851404 | 10 | 96643549 | -0.834 | 0.040 | 2.67x10^-96^ |  |
| cg07270175 | 10 | 96123049 | 0.638 | 0.031 | 3.85x10^-95^ | *NOC3L* |
| cg11265800 | 10 | 96312726 | 0.397 | 0.028 | 3.47x10^-45^ | *HELLS* |
| cg09036531 | 10 | 96991505 | -0.602 | 0.043 | 4.23x10^-45^ |  |
| ch.10.1999173F | 10 | 95990357 | 0.297 | 0.024 | 1.55x10^-34^ | *PLCE1* |
| cg11380830 | 10 | 96123085 | -0.238 | 0.023 | 8.54x10^-26^ | *NOC3L* |
| cg11119406 | 10 | 96881724 | 0.663 | 0.063 | 1.00x10^-25^ |  |
| cg27587826 | 10 | 96166801 | 0.270 | 0.027 | 2.84x10^-24^ | *TBC1D12* |
| cg14219693 | 10 | 96928076 | 0.589 | 0.060 | 1.58x10^-22^ |  |
| cg20276630 | 10 | 97055439 | 0.402 | 0.042 | 7.25x10^-22^ |  |
| cg18130464 | 10 | 96039752 | -0.308 | 0.035 | 2.48x10^-18^ | *PLCE1* |
| cg14229173 | 10 | 95974688 | -0.460 | 0.053 | 5.73x10^-18^ | *PLCE1* |
| cg26400074 | 10 | 96996533 | 0.435 | 0.050 | 6.20x10^-18^ |  |
| cg02338345 | 10 | 97036527 | -0.259 | 0.030 | 6.80x10^-18^ | *PDLIM1* |
| cg24338920 | 10 | 96075700 | 0.326 | 0.039 | 3.83x10^-17^ | *PLCE1* |
| cg11911874 | 10 | 97051104 | -0.386 | 0.046 | 9.44x10^-17^ | *PDLIM1* |
| cg03178678 | 10 | 97051225 | -0.275 | 0.035 | 4.28x10^-15^ | *PDLIM1* |
| cg15454820 | 10 | 96990858 | -0.098 | 0.013 | 1.68x10^-14^ |  |
| cg03269218 | 10 | 96990700 | -0.122 | 0.016 | 4.24x10^-14^ |  |
| cg04125153 | 10 | 95987486 | -0.481 | 0.064 | 7.53x10^-14^ | *PLCE1* |
| cg19476376 | 10 | 96990921 | -0.120 | 0.017 | 4.75x10^-13^ |  |
| cg04539301 | 10 | 96990923 | -0.138 | 0.019 | 6.71x10^-13^ |  |
| cg07347315 | 10 | 97003301 | 0.164 | 0.023 | 9.88x10^-13^ | *PDLIM1* |
| cg24087710 | 10 | 96928657 | 0.341 | 0.050 | 1.24x10^-11^ |  |
| cg15776783 | 10 | 96974685 | -0.255 | 0.038 | 1.61x10^-11^ | *ACSM6* |
| cg05599883 | 10 | 97051319 | -0.219 | 0.033 | 2.13x10^-11^ | *PDLIM1* |
| cg15233961 | 10 | 96990543 | -0.148 | 0.022 | 2.18x10^-11^ |  |
| cg18389639 | 10 | 97049610 | -0.383 | 0.059 | 1.09x10^-10^ | *PDLIM1* |
| cg23153757 | 12 | 33048710 | -0.192 | 0.030 | 1.24x10^-10^ | *PKP2* |
| cg01529847 | 10 | 96161365 | -0.286 | 0.046 | 6.09x10^-10^ | *TBC1D12* |
| cg04508033 | 10 | 96047603 | 0.210 | 0.035 | 1.87x10^-09^ | *PLCE1* |
| cg02989450 | 10 | 96104839 | -0.103 | 0.017 | 3.32x10^-09^ | *NOC3L* |
| cg14196507 | 10 | 96443383 | 0.232 | 0.039 | 3.83x10^-09^ | *CYP2C18* |
| cg25841553 | 10 | 96356520 | -0.111 | 0.019 | 4.70x10^-09^ | *HELLS* |
| cg10164249 | 10 | 96121853 | 0.095 | 0.016 | 8.55x10^-09^ | *NOC3L* |
| cg06570967 | 10 | 96989650 | -0.214 | 0.038 | 1.61x10^-08^ |  |
| cg13435317 | 10 | 95826508 | -0.118 | 0.021 | 3.62x10^-08^ | *PLCE1* |
| cg27423310 | 10 | 97051091 | -0.185 | 0.034 | 4.13x10^-08^ | *PDLIM1* |
| cg10304160 | 10 | 95457300 | 0.156 | 0.029 | 6.58x10^-08^ | *FRA10AC1* |

CHR: chromosome; BP: base pair; SE: standard error

Table S2 Significant CpGs associated with the linear term of CYP2C19 metaboliser status (MOA MWAS analysis).

| CpG | CHR | BP | B | SE | P | Nearest gene |
| --- | --- | --- | --- | --- | --- | --- |
| cg20031717 | 10 | 96523248 | -1.626 | 0.040 | <2.22x10^-308^ | *CYP2C19* |
| cg02808805 | 10 | 96521820 | 1.231 | 0.032 | <2.22x10^-308^ | *CYP2C19* |
| cg07889765 | 10 | 96123159 | -0.802 | 0.021 | <2.22x10^-308^ | *NOC3L* |
| cg10751070 | 10 | 96143568 | 1.397 | 0.025 | <2.22x10^-308^ |  |
| cg11776334 | 10 | 96046836 | -1.515 | 0.042 | 1.76x10^-291^ | *PLCE1* |
| cg08923894 | 10 | 96123172 | -0.661 | 0.019 | 1.01x10^-277^ | *NOC3L* |
| cg23961982 | 10 | 96109291 | -1.230 | 0.036 | 2.84x10^-255^ | *NOC3L* |
| cg16964198 | 10 | 96199371 | -0.961 | 0.029 | 1.09x10^-246^ | *TBC1D12* |
| cg00051662 | 10 | 96521086 | 0.827 | 0.030 | 1.59x10^-169^ | *CYP2C19* |
| cg15851404 | 10 | 96643549 | -0.834 | 0.040 | 1.70x10^-97^ |  |
| cg07270175 | 10 | 96123049 | 0.638 | 0.031 | 2.38x10^-96^ | *NOC3L* |
| cg11265800 | 10 | 96312726 | 0.397 | 0.028 | 2.58x10^-46^ | *HELLS* |
| cg09036531 | 10 | 96991505 | -0.602 | 0.042 | 3.15x10^-46^ |  |
| ch.10.1999173F | 10 | 95990357 | 0.297 | 0.024 | 1.75x10^-35^ | *PLCE1* |
| cg11380830 | 10 | 96123085 | -0.238 | 0.022 | 1.51x10^-26^ | *NOC3L* |
| cg11119406 | 10 | 96881724 | 0.663 | 0.062 | 1.78x10^-26^ |  |
| cg27587826 | 10 | 96166801 | 0.270 | 0.026 | 5.49x10^-25^ | *TBC1D12* |
| cg14219693 | 10 | 96928076 | 0.589 | 0.059 | 3.39x10^-23^ |  |
| cg20276630 | 10 | 97055439 | 0.402 | 0.041 | 1.61x10^-22^ |  |
| cg18130464 | 10 | 96039752 | -0.308 | 0.035 | 6.91x10^-19^ | *PLCE1* |
| cg14229173 | 10 | 95974688 | -0.460 | 0.052 | 1.63x10^-18^ | *PLCE1* |
| cg26400074 | 10 | 96996533 | 0.435 | 0.050 | 1.77x10^-18^ |  |
| cg02338345 | 10 | 97036527 | -0.259 | 0.030 | 1.95x10^-18^ | *PDLIM1* |
| cg24338920 | 10 | 96075700 | 0.326 | 0.038 | 1.15x10^-17^ | *PLCE1* |
| cg11911874 | 10 | 97051104 | -0.386 | 0.046 | 2.92x10^-17^ | *PDLIM1* |
| cg03178678 | 10 | 97051225 | -0.275 | 0.034 | 1.48x10^-15^ | *PDLIM1* |
| cg15454820 | 10 | 96990858 | -0.098 | 0.013 | 6.05x10^-15^ |  |
| cg03269218 | 10 | 96990700 | -0.122 | 0.016 | 1.57x10^-14^ |  |
| cg04125153 | 10 | 95987486 | -0.481 | 0.063 | 2.84x10^-14^ | *PLCE1* |
| cg19476376 | 10 | 96990921 | -0.120 | 0.016 | 1.90x10^-13^ |  |
| cg04539301 | 10 | 96990923 | -0.138 | 0.019 | 2.71x10^-13^ |  |
| cg07347315 | 10 | 97003301 | 0.164 | 0.023 | 4.03x10^-13^ | *PDLIM1* |
| cg24087710 | 10 | 96928657 | 0.341 | 0.049 | 5.48x10^-12^ |  |
| cg15776783 | 10 | 96974685 | -0.255 | 0.037 | 7.16x10^-12^ | *ACSM6* |
| cg05599883 | 10 | 97051319 | -0.219 | 0.032 | 9.55x10^-12^ | *PDLIM1* |
| cg15233961 | 10 | 96990543 | -0.148 | 0.022 | 9.79x10^-12^ |  |
| cg18389639 | 10 | 97049610 | -0.383 | 0.058 | 5.16x10^-11^ | *PDLIM1* |
| cg23153757 | 12 | 33048710 | -0.192 | 0.029 | 5.88x10^-11^ | *PKP2* |
| cg01529847 | 10 | 96161365 | -0.286 | 0.045 | 3.05x10^-10^ | *TBC1D12* |
| cg04508033 | 10 | 96047603 | 0.210 | 0.034 | 9.72x10^-10^ | *PLCE1* |
| cg02989450 | 10 | 96104839 | -0.103 | 0.017 | 1.75x10^-09^ | *NOC3L* |
| cg14196507 | 10 | 96443383 | 0.232 | 0.039 | 2.03x10^-09^ | *CYP2C18* |
| cg25841553 | 10 | 96356520 | -0.111 | 0.019 | 2.51x10^-09^ | *HELLS* |
| cg10164249 | 10 | 96121853 | 0.095 | 0.016 | 4.66x10^-09^ | *NOC3L* |
| cg06570967 | 10 | 96989650 | -0.214 | 0.037 | 8.97x10^-09^ |  |
| cg13435317 | 10 | 95826508 | -0.118 | 0.021 | 2.07x10^-08^ | *PLCE1* |
| cg27423310 | 10 | 97051091 | -0.185 | 0.033 | 2.37x10^-08^ | *PDLIM1* |
| cg10304160 | 10 | 95457300 | 0.156 | 0.028 | 3.83x10^-08^ | *FRA10AC1* |

CHR: chromosome; BP: base pair; SE: standard error

Table S3 Significant CpGs associated with the quadratic term of CYP2C19 metaboliser status

| CpG | CHR | BP | B | SE | P | Nearest gene |
| --- | --- | --- | --- | --- | --- | --- |
| cg10751070 | 10 | 96143568 | 1.478 | 0.028 | <2.22x10^-308^ |  |
| cg16964198 | 10 | 96199371 | 2.325 | 0.028 | <2.22x10^-308^ | *TBC1D12* |
| cg20031717 | 10 | 96523248 | 1.775 | 0.044 | <2.22x10^-308^ | *CYP2C19* |
| **cg08280358** | 10 | 96189867 | 0.919 | 0.036 | 1.04x10^-143^ | *TBC1D12* |
| cg08923894 | 10 | 96123172 | -0.478 | 0.021 | 1.47x10^-114^ | *NOC3L* |
| cg07889765 | 10 | 96123159 | -0.551 | 0.024 | 2.53x10^-114^ | *NOC3L* |
| **cg08925046** | 10 | 97008920 | -1.420 | 0.065 | 2.97x10^-106^ | *PDLIM1* |
| cg14219693 | 10 | 96928076 | -1.400 | 0.065 | 1.56x10^-102^ |  |
| cg11776334 | 10 | 96046836 | -0.906 | 0.047 | 6.35x10^-82^ | *PLCE1* |
| cg24087710 | 10 | 96928657 | -1.033 | 0.054 | 6.54x10^-81^ |  |
| cg23961982 | 10 | 96109291 | -0.740 | 0.041 | 3.47x10^-73^ | *NOC3L* |
| cg02808805 | 10 | 96521820 | -0.636 | 0.036 | 3.93x10^-68^ | *CYP2C19* |
| cg15851404 | 10 | 96643549 | 0.602 | 0.044 | 8.19x10^-42^ |  |
| cg00051662 | 10 | 96521086 | -0.413 | 0.034 | 1.06x10^-34^ | *CYP2C19* |
| cg26400074 | 10 | 96996533 | 0.666 | 0.055 | 6.29x10^-34^ |  |
| cg09036531 | 10 | 96991505 | -0.510 | 0.047 | 1.24x10^-27^ |  |
| **cg21800396** | 10 | 96968197 | -0.476 | 0.048 | 6.95x10^-23^ | *ACSM6* |
| **cg08883204** | 10 | 97069028 | 0.474 | 0.050 | 1.14x10^-21^ |  |
| cg07270175 | 10 | 96123049 | 0.317 | 0.034 | 2.36x10^-20^ | *NOC3L* |
| cg04539301 | 10 | 96990923 | -0.193 | 0.021 | 2.49x10^-20^ |  |
| cg11380830 | 10 | 96123085 | -0.225 | 0.025 | 1.08x10^-19^ | *NOC3L* |
| cg19476376 | 10 | 96990921 | -0.161 | 0.018 | 3.43x10^-19^ |  |
| **cg17725512** | 10 | 96447808 | 0.206 | 0.023 | 6.85x10^-19^ | *CYP2C18* |
| cg02338345 | 10 | 97036527 | -0.287 | 0.033 | 1.69x10^-18^ | *PDLIM1* |
| cg18389639 | 10 | 97049610 | -0.562 | 0.065 | 3.16x10^-18^ | *PDLIM1* |
| cg15233961 | 10 | 96990543 | -0.208 | 0.024 | 4.87x10^-18^ |  |
| **cg00087741** | 10 | 96961488 | -0.511 | 0.059 | 5.33x10^-18^ | *ACSM6* |
| cg15454820 | 10 | 96990858 | -0.120 | 0.014 | 6.99x10^-18^ |  |
| **cg13512927** | 10 | 95984834 | 0.447 | 0.052 | 1.18x10^-17^ | *PLCE1* |
| cg24338920 | 10 | 96075700 | 0.352 | 0.042 | 7.15x10^-17^ | *PLCE1* |
| **cg14302996** | 10 | 97205147 | 0.514 | 0.063 | 3.36x10^-16^ | *SORBS1* |
| **cg20426415** | 10 | 96446783 | 0.301 | 0.038 | 1.88x10^-15^ | *CYP2C18* |
| cg03269218 | 10 | 96990700 | -0.139 | 0.018 | 3.07x10^-15^ |  |
| **cg17014018** | 10 | 96442621 | 0.338 | 0.044 | 2.90x10^-14^ | *CYP2C18* |
| cg06570967 | 10 | 96989650 | -0.311 | 0.041 | 4.68x10^-14^ |  |
| cg11119406 | 10 | 96881724 | 0.508 | 0.069 | 1.92x10^-13^ |  |
| **cg12575696** | 10 | 96998139 | -0.444 | 0.064 | 5.94x10^-12^ | *PDLIM1* |
| cg11265800 | 10 | 96312726 | -0.212 | 0.031 | 6.34x10^-12^ | *HELLS* |
| **cg21636366** | 10 | 96447855 | -0.195 | 0.030 | 7.97x10^-11^ | *CYP2C18* |
| **cg25674102** | 10 | 24684432 | 0.506 | 0.079 | 1.55x10^-10^ | *KIAA1217* |
| **cg23210118** | 10 | 95653378 | -0.161 | 0.026 | 3.98x10^-10^ | *TMEM20* |
| **cg15160630** | 10 | 96048116 | 0.422 | 0.069 | 8.45x10^-10^ | *PLCE1* |
| **cg03013070** | 10 | 95965579 | -0.347 | 0.057 | 9.04x10^-10^ | *PLCE1* |
| **cg04708601** | 6 | 101880078 | 0.280 | 0.048 | 5.39x10^-09^ | *GRIK2* |
| cg07347315 | 10 | 97003301 | 0.145 | 0.025 | 7.12x10^-09^ | *PDLIM1* |
| ch.10.1999173F | 10 | 95990357 | 0.153 | 0.027 | 7.97x10^-09^ | *PLCE1* |
| **cg09847250** | 16 | 65397909 | -0.281 | 0.051 | 2.80x10^-08^ | *LOC283867* |
| **cg14013452** | 10 | 96130481 | -0.203 | 0.037 | 4.91x10^-08^ |  |

CHR: chromosome; BP: base pair; SE: standard error

Bold CpG sites showed non-linear associations with CYP2C19 metaboliser status

Table S4 Significance of the interaction effects of the CYP2C19 metaboliser status and CYP2C19-metabolised medication use on M-values of significant non-linear CpG sites

| CpG | P_interaction_ | P_FDR_ |
| --- | --- | --- |
| cg08280358 | 0.405 | 0.640 |
| cg08925046 | 0.021 | 0.147 |
| cg21800396 | 0.059 | 0.226 |
| cg08883204 | 0.399 | 0.640 |
| cg17725512 | 0.025 | 0.147 |
| cg00087741 | 0.526 | 0.714 |
| cg13512927 | 0.671 | 0.750 |
| cg14302996 | 0.031 | 0.147 |
| cg20426415 | 0.665 | 0.750 |
| cg17014018 | 0.455 | 0.665 |
| cg12575696 | 0.169 | 0.458 |
| cg21636366 | 0.214 | 0.509 |
| cg25674102 | 0.331 | 0.628 |
| cg23210118 | 0.773 | 0.773 |
| cg15160630 | 0.285 | 0.601 |
| cg03013070 | 0.118 | 0.375 |
| cg04708601 | 0.014 | 0.147 |
| cg09847250 | 0.743 | 0.773 |
| cg14013452 | 0.588 | 0.745 |

P_interaction_: P values for the interaction term of CYP2C19 metaboliser status (categorised as three groups: normal, rapid/intermediate, and ultrarapid/poor) and CYP2C19-related medication use (categorised as four groups: not current user, inducer, inhibitor, and substrate); P_FDR_: P values for interaction term after FDR correction

Table S5 Significant CpG sites detected from linear model MWAS analysis after additional adjustment for M-value of cg20031717.

| CpG | CHR | BP | B | SE | P | Nearest gene |
| --- | --- | --- | --- | --- | --- | --- |
| cg07889765 | 10 | 96123159 | -0.776 | 0.020 | <2.22x10^-308^ | *NOC3L* |
| cg10751070 | 10 | 96143568 | 1.404 | 0.022 | <2.22x10^-308^ |  |
| cg11776334 | 10 | 96046836 | -1.461 | 0.039 | 5.15x10^-310^ | *PLCE1* |
| cg08923894 | 10 | 96123172 | -0.643 | 0.017 | 5.89x10^-300^ | *NOC3L* |
| cg23961982 | 10 | 96109291 | -1.210 | 0.034 | 7.24x10^-281^ | *NOC3L* |
| cg02808805 | 10 | 96521820 | 1.059 | 0.030 | 5.35x10^-269^ | *CYP2C19* |
| cg00051662 | 10 | 96521086 | 0.695 | 0.029 | 7.95x10^-130^ | *CYP2C19* |
| cg16964198 | 10 | 96199371 | -0.659 | 0.029 | 3.48x10^-115^ | *TBC1D12* |
| cg07270175 | 10 | 96123049 | 0.636 | 0.029 | 3.95x10^-104^ | *NOC3L* |
| cg15851404 | 10 | 96643549 | -0.767 | 0.038 | 1.79x10^-89^ |  |
| cg09036531 | 10 | 96991505 | -0.636 | 0.041 | 3.81x10^-55^ |  |
| cg11265800 | 10 | 96312726 | 0.330 | 0.027 | 1.89x10^-34^ | *HELLS* |
| ch.10.1999173F | 10 | 95990357 | 0.275 | 0.023 | 1.46x10^-32^ | *PLCE1* |
| cg26400074 | 10 | 96996533 | 0.514 | 0.048 | 1.14x10^-26^ |  |
| cg11119406 | 10 | 96881724 | 0.636 | 0.060 | 5.27x10^-26^ |  |
| cg27587826 | 10 | 96166801 | 0.260 | 0.025 | 9.80x10^-25^ | *TBC1D12* |
| cg11380830 | 10 | 96123085 | -0.207 | 0.022 | 1.06x10^-21^ | *NOC3L* |
| cg14229173 | 10 | 95974688 | -0.482 | 0.051 | 2.34x10^-21^ | *PLCE1* |
| cg24338920 | 10 | 96075700 | 0.346 | 0.037 | 7.37x10^-21^ | *PLCE1* |
| cg15454820 | 10 | 96990858 | -0.113 | 0.012 | 1.49x10^-20^ |  |
| cg03269218 | 10 | 96990700 | -0.140 | 0.015 | 8.39x10^-20^ |  |
| cg02338345 | 10 | 97036527 | -0.256 | 0.029 | 3.94x10^-19^ | *PDLIM1* |
| cg14219693 | 10 | 96928076 | 0.510 | 0.058 | 8.03x10^-19^ |  |
| cg19476376 | 10 | 96990921 | -0.138 | 0.016 | 1.64x10^-18^ |  |
| cg18130464 | 10 | 96039752 | -0.291 | 0.034 | 4.42x10^-18^ | *PLCE1* |
| cg04539301 | 10 | 96990923 | -0.159 | 0.018 | 4.99x10^-18^ |  |
| cg15233961 | 10 | 96990543 | -0.171 | 0.021 | 4.87x10^-16^ |  |
| cg20276630 | 10 | 97055439 | 0.313 | 0.040 | 4.89x10^-15^ |  |
| cg18389639 | 10 | 97049610 | -0.442 | 0.057 | 6.18x10^-15^ | *PDLIM1* |
| cg11911874 | 10 | 97051104 | -0.341 | 0.044 | 1.31x10^-14^ | *PDLIM1* |
| cg03178678 | 10 | 97051225 | -0.257 | 0.033 | 1.45x10^-14^ | *PDLIM1* |
| cg07347315 | 10 | 97003301 | 0.168 | 0.022 | 2.16x10^-14^ | *PDLIM1* |
| cg05599883 | 10 | 97051319 | -0.230 | 0.031 | 1.69x10^-13^ | *PDLIM1* |
| cg06570967 | 10 | 96989650 | -0.225 | 0.036 | 4.69x10^-10^ |  |
| cg04125153 | 10 | 95987486 | -0.382 | 0.061 | 4.93x10^-10^ | *PLCE1* |
| cg05771722 | 10 | 95766961 | -0.275 | 0.045 | 7.94x10^-10^ | *PLCE1* |
| cg04508033 | 10 | 96047603 | 0.204 | 0.033 | 9.05x10^-10^ | *PLCE1* |
| cg00087741 | 10 | 96961488 | -0.307 | 0.052 | 3.40x10^-09^ | *ACSM6* |
| cg13435317 | 10 | 95826508 | -0.118 | 0.020 | 7.72x10^-09^ | *PLCE1* |
| cg15776783 | 10 | 96974685 | -0.207 | 0.036 | 9.24x10^-09^ | *ACSM6* |
| cg25841553 | 10 | 96356520 | -0.102 | 0.018 | 1.94x10^-08^ | *HELLS* |
| cg10164249 | 10 | 96121853 | 0.086 | 0.016 | 4.64x10^-08^ | *NOC3L* |
| cg23153757 | 12 | 33048710 | -0.154 | 0.028 | 6.53x10^-08^ | *PKP2* |

CHR: chromosome; BP: base pair; SE: standard error
